# Linguistic processing of task-irrelevant speech at a Cocktail Party

**DOI:** 10.1101/2020.11.08.373746

**Authors:** Paz Har-shai Yahav, Elana Zion Golumbic

**Affiliations:** The Gonda Center for Multidisciplinary Brain Research, Bar Ilan University, Israel

## Abstract

Paying attention to one speaker in noisy environments can be extremely difficult, because to-be-attended and task-irrelevant speech compete for processing resources. We tested whether this competition is restricted to acoustic-phonetic interference or if it extends to competition for linguistic processing as well. Neural activity was recorded using Magnetoencephalography as human participants were instructed to attended to natural speech presented to one ear, and task-irrelevant stimuli were presented to the other. Task-irrelevant stimuli consisted either of random sequences of syllables, or syllables structured to form coherent sentences, using hierarchical frequency-tagging.

We find that the phrasal structure of structured task-irrelevant stimuli was represented in the neural response in left inferior frontal and posterior parietal regions, indicating that selective attention does not fully eliminate linguistic processing of task-irrelevant speech. Additionally, neural tracking of to-be-attended speech in left inferior frontal regions was enhanced when competing with structured task-irrelevant stimuli, suggesting inherent competition between them for linguistic processing.

**Impact Statement:** Syntactic structure-building processes can be applied to speech that is task-irrelevant and should be ignored, demonstrating that Selective Attention does not fully eliminate linguistic processing of competing speech.

## Introduction

The seminal speech-shadowing experiments conducted in the 50s and 60s set the stage for studying one of the primary cognitive challenges encountered in daily life: how do our perceptual and linguistic systems deal effectively with competing speech inputs? (Cherry 1953; Broadbent 1958; Treisman 1960). Over the past decades, a wealth of behavioral and neural evidence has accumulated showing that when only one speech-stream is behaviorally relevant, auditory and linguistic resources are devoted to its preferential encoding at the expense of other task-irrelevant input. Consequentially, this so-called ‘attended’ message can be repeated, comprehended, and remembered, whereas very little of competing task-irrelevant speech is explicitly recalled (Glucksberg and Cowen 1970; Ambler et al. 1976; Neely and LeCompte 1999; Oswald et al. 2000; Brungart et al. 2001). This attentional selection is accompanied by attenuation of speech-tracking for unattended speech in auditory regions (Mesgarani and Chang 2012; Horton et al. 2013; Zion Golumbic, Cogan, et al. 2013; O’Sullivan et al. 2015; Fiedler et al. 2019; Teoh and Lalor 2019) as well as language-related regions (Zion Golumbic, Ding, et al. 2013), particularly for linguistic-features of the speech (Brodbeck, Hong, et al. 2018; Broderick et al. 2018; Ding et al. 2018; Brodbeck et al. 2020a). However, as demonstrated even in the earliest studies, the content of task-irrelevant speech is probably not fully suppressed and can affect listener behavior in a variety of ways (Moray 1959; Bryden 1964; Yates 1965). Indeed, despite decades of research, the extent to which concurrent speech are processed and the nature of the competition for resources between ‘attended’ and ‘task-irrelevant’ input in multi-speaker contexts is still highly debated (Kahneman 1973; Driver 2001; Lachter et al. 2004; Bronkhorst 2015).

Fueling this debate are often-conflicting empirical findings regarding whether or not task-irrelevant speech is processed for semantic and linguistic content. Many studies fail to find behavioral or neural evidence for processing task-irrelevant speech beyond its acoustic features (Carlyon et al. 2001; Lachter et al. 2004; Ding et al. 2018). However, others are able to demonstrate that at least some linguistic information is gleaned from task-irrelevant speech. For example, task-irrelevant speech is more distracting than non-speech or incomprehensible distractors (Rhebergen et al. 2005; Iyer et al. 2010; Best et al. 2012; Gallun and Diedesch 2013; Carey et al. 2014; Kilman et al. 2014; Swaminathan et al. 2015; Kidd et al. 2016), and there are also indications for implicit processing of the semantic content of task-irrelevant speech, manifest through priming effects or memory intrusions (Tun et al. 2002; Dupoux et al. 2003; Rivenez et al. 2006; Beaman et al. 2007; Carey et al. 2014; Aydelott et al. 2015; Schepman et al. 2016). Known as the ‘Irrelevant Sound Effect’ (ISE), these are not always accompanied by explicit recall or recognition (Lewis 1970; Bentin et al. 1995; Röer et al. 2017a), although in some cases task-irrelevant words, such as one’s own name, may also ‘break in consciousness’ (Cherry 1953; Treisman 1960; Wood and Cowan 1995; Conway et al. 2001).

Behavioral findings indicating linguistic processing of task-irrelevant speech have been interpreted in two opposing ways. Proponents of Late-Selection attention theories understand them as reflecting the system’s capability to apply linguistic processing to more than one speech stream in parallel, albeit mostly pre-consciously (Deutsch and Deutsch 1963; Parmentier 2008; Parmentier et al. 2018; Vachon et al. 2019). However, others maintain an Early-Selection perspective, namely, that only one speech stream can be processed linguistically due to inherent processing bottlenecks, but that listeners may shift their attention between concurrent streams giving rise to occasional (conscious or pre-conscious) intrusions from task-irrelevant speech (Cooke 2006; Vestergaard et al. 2011; Fogerty et al. 2018). Adjudicating between these two explanations experimentally is difficult, due to the largely indirect-nature of the operationalizations used to assess linguistic processing of task-irrelevant speech. Moreover, much of the empirical evidence fueling this debate focuses on detection of individual ‘task-irrelevant’ words, effects that can be easily explained either by parallel processing or by attention-shifts, due to their short duration.

In attempt to broaden this conversation, here we use objective neural measures to evaluate the level of processing applied to task-irrelevant speech. Using a previously established technique of hierarchical frequency-tagging (Ding et al. 2016; Makov et al. 2017), we are able to go beyond the question of detecting individual words and probe whether linguistic processes that require integration over longer periods of time – such as syntactic structure building – are applied to task-irrelevant speech. To study this, we recorded brain activity using Magnetoencephalography (MEG) during a dichotic listening selective-attention experiment. Participants were instructed to attend to narratives of natural speech presented to one ear, and to ignore speech input from the other ear (Figure 1). Task-irrelevant stimuli consisted of sequences of syllables, presented at a constant rate (4Hz), with their order manipulated to either create linguistically Structured or Non-Structured sequences. Specifically, for the Non-Structured stimuli syllables were presented in a completely random order, whereas in the Structured stimuli syllables are ordered to form coherent words, phrases and sentences. In keeping with the frequency-tagging approach, each of these linguistic levels is associated with a different frequency (words - 2Hz, phrases - 1Hz, sentences - 0.5Hz). By structuring task-irrelevant speech in this way, the two conditions were perfectly controlled for low-level acoustic attributes that contribute to energetic masking (e.g. loudness, pitch, and fine-structure), as well as for the presence of recognizable acoustic-phonetic units, which proposedly contributes to phonetic interference during speech-on-speech masking (Rhebergen et al. 2005; Shinn-Cunningham 2008; Kidd et al. 2016). Rather, the only difference between the conditions was in the order of the syllables which either did or did not form linguistic structures. Consequentially, if the neural signal shows peaks at frequencies associated with linguistic-features of Structured task-irrelevant speech, as has been reported previously when these type of stimuli are attended or presented without competition (Ding et al. 2016, 2018; Makov et al. 2017), this would provide evidence that integration-based processes operating on longer time-scales are applied to task-irrelevant speech, for identifying longer linguistic units comprised of several syllables. In addition, we also tested whether the neural encoding of the to-be-attended speech itself was affected by the linguistic structure of task-irrelevant speech, which could highlight the source of potential tradeoffs or competition for resources when presented with competing speech (Zion Golumbic, Ding, et al. 2013; O’Sullivan et al. 2015; Fiedler et al. 2019; Teoh and Lalor 2019).

**Figure 1.**
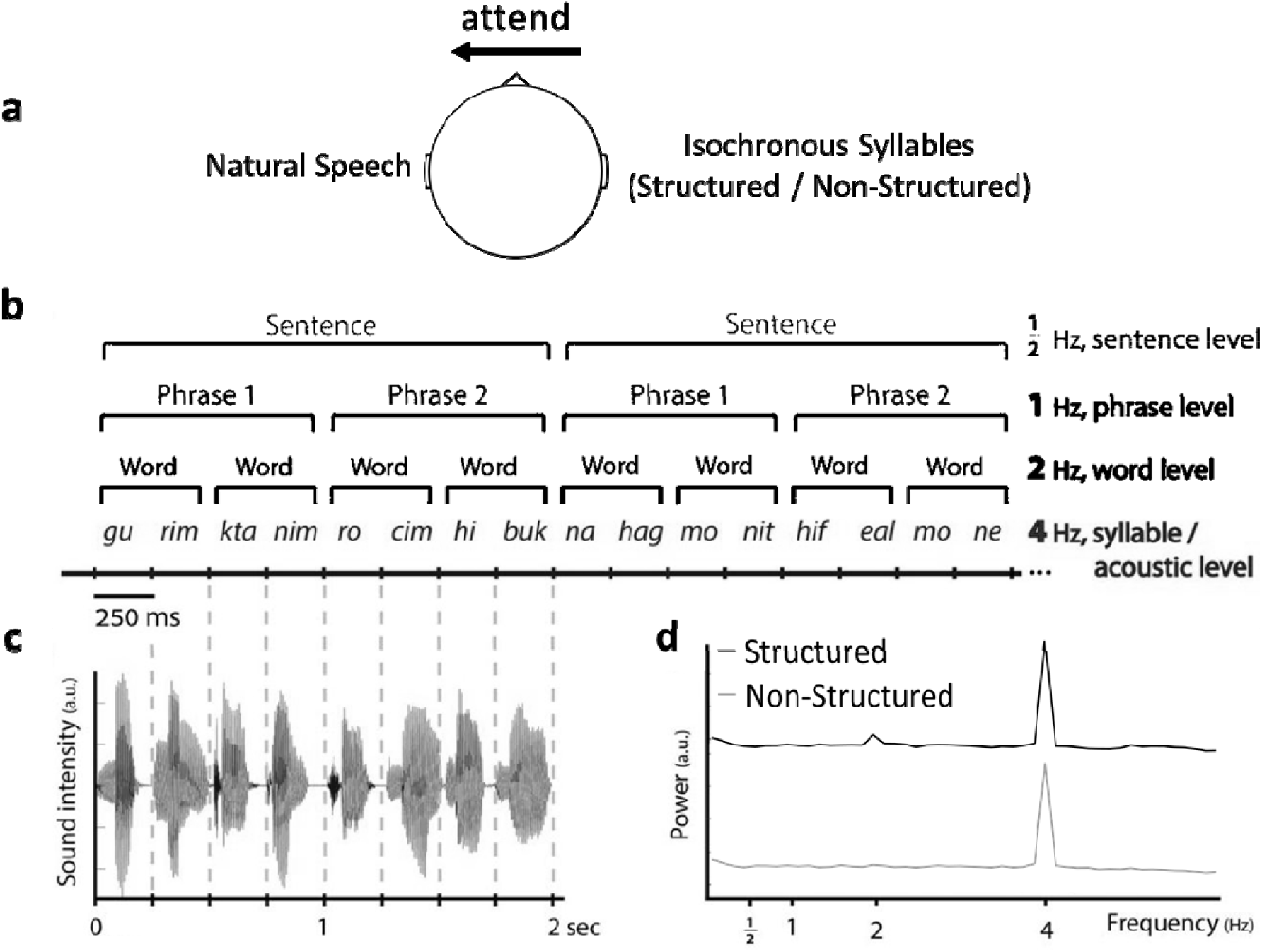
**a**, Dichotic paradigm experiment. Participants were instructed to attended right or left ear (counterbalanced) and ignore the other. To-be-attended stimuli was always natural Hebrew speech, with multiple answers questions about the content at the end of each trial. The task-irrelevant ear was always presented with hierarchical frequency-tagging stimuli in two conditions: Structured and Non-Structured. **b**, Example of intelligible (Structured) speech composed of 250 ms syllables in which 4 levels of information are differentiated based on their rate: acoustic/syllabic, word, phrasal, and sentential rates (at 4, 2, 1, and 0.5Hz, respectively). Translation of the Hebrew sentences in the example: “Small puppies want a hug” and “A taxi driver turned on the meter.” Control stimuli were unintelligible (Non-Structured) syllable sequences with similar syllabic-rate of 4Hz. **c**, Representative sound wave of a single sentence (2s). Sound intensity fluctuates at the rate of 4Hz. **d**, Induced power spectrum of 50 stimuli soundwave envelope per condition. * Figure adapted from Makov et al. 2017.

## Materials and Methods

### Participants

We measured MEG recordings from 30 (18 females, 12 males) native Hebrew speakers. Participants were adult volunteers, ages ranging between 18 and 34 (M=24.8, SD=±4.2), and all were right-handed. Sample size was determined a-priori, based on a previous study from our group using a similar paradigm and electrophysiological measures (Makov et al. 2017), where significant effects were found in a sample of n=21 participants. Exclusion criteria for participation included: non-native Hebrew speakers, a history of neurological disorders or ADHD (based on self-report) or the existence of metal implants (which would disrupt MEG recordings). The study was approved by the IRB committee at Bar-Ilan University and all participants provided their written consent for participation prior to the experiment.

### Natural Speech (to-be-attended)

Natural speech stimuli were narratives from publicly available Hebrew podcasts and short audio stories (duration: 44.53±3.23 seconds). These speech materials were chosen from an existing database in the lab, that were used in previous studies and for which the behavioral task had already been validated (see Experimental Procedure). The stimuli originally consisted of narratives in both female and male voices. However, since it is known that selective attention to speech is highly influenced by whether the competing voices are of the same/different sex (Brungart et al. 2001; Rivenez et al. 2006; Ding and Simon 2012), and since the task-irrelevant stimuli were recorded only in a male voice (see below), we transformed narratives that were originally recorded in a female voice to a male voice (change-gender function in Praat; Boersma P. 2011, www.praat.org). To ensure that the gender change did not affect the naturalness of the speech and to check for abnormalities in the materials, we conducted a short survey among 10 native Hebrew speakers. They all agreed that the speech sounded natural and normal. Sound intensity was equated across all narratives. Stimuli examples are available at: https://osf.io/e93qa. These natural speech narratives served as the to-be-attended stimuli in the experiment. For each participant they were randomly paired with task-irrelevant speech (regardless of condition), to avoid material-specific effects.

### Frequency-tagged Speech (task-irrelevant)

A bank of individually recorded Hebrew syllables were used to create two sets of isochronous speech sequences. Single syllables were recorded in random order by a male actor, and remaining prosodic cues were removed using pitch normalization in Praat. Additional sound editing was performed to adjust the length of each syllable to be precisely 250 ms either by truncation or silence padding at the end (original mean duration 243.6±64.3 ms, range 168–397 ms). In case of truncation, a fading out effect was applied to the last 25 ms to avoid clicks. Sound intensity was then manually equated for all syllables.

These syllables were concatenated to create long sequences using custom-written scripts in MATLAB (The MathWorks), equated in length to those of the natural speech segments (44.53±3.23). Sequences could either be Non-Structured, with syllables presented in a fully random order without creating meaningful linguistic units, or they could be linguistically Structured. Structured sequences were identical to those used in a previous study from our group (Makov et al. 2017), and were formed as follows: Every two syllables formed a word, every two words formed a phrase, and every two phrases formed a sentence. Because syllables were grouped hierarchically into linguistic constituents with no additional acoustic gaps inserted between them, different linguistic hierarchies are associated with fixed periodicities throughout the stimuli (syllables at 4Hz, words at 2Hz, phrases at 1Hz, and sentences at 0.5Hz; Figure 1b). Structured stimuli also contained no prosodic cues or other low-level acoustic indications for boundaries between linguistics structures, nor did Structured sentences include rhymes, passive form of verbs, or arousing semantic content. See Supplementary Material for more information on the construction of Structured and Non-Structured stimuli.

The modulation spectrum of both types of task-irrelevant stimuli is shown in Figure 1d. It was calculated using a procedure analogous to the spectral analysis performed on the MEG data, in order to ensure maximal comparability between the spectrum of the stimuli and the spectrum of the neural response. Specifically, (1) the broadband envelope of each sequence was extracted by taking the root-mean-square of the audio (10-ms smoothing window); (2) the envelope was segmented into 8-second long segments, which was identical to the segmentation of the MEG data; (3) a fast Fourier transform (FFT) was applied to each segment; and (4) averaged across segments. As expected, both stimuli contained a prominent peak at 4Hz, corresponding to the syllable-rate. The Structured stimuli also contain a smaller peak at 2Hz, which corresponds to the word-rate. This is an undesirable side-effect of the frequency-tagging approach, since ideally these stimuli should not contain any energy at frequencies other than the syllable-rate. As shown in the Supplementary Material, the 2Hz peak in the modulation spectrum reflects the fact that a consistently different subset of syllables occurs at each position within the sequence (e.g. at the beginning\end of words; for similar acoustic effects when using frequency-tagging of multi-syllable words see (Luo and Ding 2020). A similar 2Hz peak is not observed in the Non-Structured condition, where the syllables are randomly positioned throughout all stimuli. Given this difference in the modulation spectrum, if we were to observe a 2Hz peak in the neural response to Structured vs. Non-Structured stimuli, this would not necessarily provide conclusive evidence for linguistic ‘word-level’ encoding (although see Makov et al. 2017 and Supplementary Material for a way to control for this). As it happened, in the current dataset we did not see a 2Hz peak in the neural response in either condition (see Results), therefore this caveat did not affect the interpretability of the data in this specific instance. Importantly, neither the Structured nor the Non-Structured stimuli contained peaks at frequencies corresponding to other linguistic levels (1Hz and 0.5Hz), hence comparison of neural responses at these frequencies remained experimentally valid. Stimuli examples are available at: https://osf.io/e93qa.

### Experimental Procedure

The experiment used a dichotic listening paradigm, in which participants were instructed to attend to a narrative of natural Hebrew speech presented to one ear, and to ignore input from the other ear where either Structured or Non-Structured isochronous task-irrelevant speech was presented. The experiment included a total of 30 trials (44.53±3.23 sec long), and participants were informed at the beginning of each trial which ear to attend to, which was counterbalanced across trials. The sound intensity of isochronous task-irrelevant stimuli increased gradually during the first three seconds of each trial, to avoid inadvertent hints regrading word-boundaries, and the start of each trial was excluded from data analysis. After each trial, participants answered four multiple choice questions about the content of the narrative they were supposed to attend to (3-answers per question; chance level = 0.33). Some of the questions required recollection of specific details (e.g., “what color was her hat?”), and some addressed the ‘gist’ of the narrative, (e.g., “why was she sad?”). The average accuracy rate of each participant (% questions answered correctly) was calculated across all questions and narratives, separately for trials in the Structured and Non-Structured condition.

This task was chosen as a way to <u>motivate and guide</u> participants to direct attention towards the to-be-attended narrative and provide <u>verification</u> that indeed they listened to it. At the same time, we recognize that this task is not highly sensitive for gauging the full extent of processing the narrative for two reasons: (1) its sparse sampling of behavior (4 questions for a 45-second narrative); and (2) since accuracy is affected by additional cognitive factors besides attention such as short-term memory, engagement, and deductive reasoning. Indeed, behavioral screening of this task showed that performance was far from perfect even when participants listened to these narratives in a single-speaker context (i.e., without additional competing speech; average accuracy rate 0.83±0.08; n=10). Hence, we did not expect performance on this task to necessarily reflect participants’ internal attentional state. At the same time, this task is instrumental in guiding participants’ selective attention toward the designated speaker, allowing us to analyze their neural activity during uninterrupted listening to continuous speech, which was the primarily goal of the current study.

### Additional Tasks

Based on previous experience, the Structured speech materials are not immediately recognizable as Hebrew speech and require some familiarization. Hence, to ensure that all participants could identify these as speech, and in order to avoid any perceptual learning effects during the main experiment, they underwent a familiarization stage prior to the start of the main experiment, inside the MEG. In this stage, participants heard sequences of 8 isochronous syllables, which were either Structured – forming a single sentence – or Non-Structured. After each trial participants were asked to repeat the sequence out loud. The familiarity stage continued until participants correctly repeated five stimuli of each type. Structured and Non-Structured stimuli were presented in random order.

At the end of the experiment a short auditory localizer task was performed. The localizer included hearing tones in five different frequencies: 400Hz, 550Hz, 700Hz, 850Hz, 1000Hz, all 200 ms long. The tones were presented with random ISIs: 500ms, 700ms, 1000ms, 1200ms, 1400ms, 1600ms, 1800ms, and 2000ms. Participants listened to the tones passively and were instructed only to focus on the fixation mark in the center of the screen.

### MEG Data Acquisition

MEG recordings were conducted with a whole-head, 248-channel magnetometer array (4D Neuroimaging, Magnes 3600 WH) in a magnetically shielded room at the Electromagnetic Brain Imaging Unit, Bar-Ilan University. A series of magnetometer and gradiometer reference coils located above the signal coils were used to record and subtract environmental noise. The location of the head with respect to the sensors was determined by measuring the magnetic field produced by small currents delivered to five head coils attached to the scalp. Before the experimental session, the position of head coils was digitized in relation to three anatomical landmarks (left and right preauricular points and nasion). The data was acquired at a sample rate of 1017.3 Hz and an online 0.1 to 200 Hz band-pass filter was applied. The 50Hz power line noise fluctuations were recorded directly from the power line as well as vibrations using a set of accelerometers attached to the sensor in order to remove the artifacts on the MEG recordings.

### MEG Preprocessing

Preprocessing was performed in MATLAB (The MathWorks) using the FieldTrip toolbox (www.fieldtriptoolbox.org). Outlier trials were identified manually by visual inspection and were excluded from analysis. Using independent component analysis (ICA) we removed eye movements (EOG), heartbeat and vibrations of the building. The clean data was then segmented into 8-second long segments, which corresponds to four sentences in the Structured condition. Critically, these segments were perfectly aligned such that they all start with the onset of a syllable, which in the Structured condition will also be the onset of a sentence.

### Source Estimation

Source estimation was performed in Python (www.python.org) using the MNE-python platform (Gramfort 2013; Gramfort et al. 2014). Source modeling was performed on the pre-processed MEG data, by computing Minimum-Norm Estimates (MNEs). In order to calculate the forward solution, and constrain source locations to the cortical surface, we constructed a Boundary Element Model (BEM) for each participant. BEM was calculated using the participants’ head shape and location relative to the MEG sensors, which was co-registered to an MRI template (FreeSurfer; surfer.nmr.mgh.harvard.edu). Then, the cortical surface of each participant was decimated to 8,194 source locations per hemisphere with at least 5-mm spacing between adjacent locations. A noise covariance matrix was estimated using the inter-trial intervals in the localizer task (see Additional Tasks), that is, periods when no auditory stimuli were presented. Then, an inverse operator was computed based on the forward solution and the noise covariance matrix, and was used to estimate the activity at each source location. For visualizing the current estimates on the cortical surface, we used dynamic Statistical Parametric Map (dSPM), which is an F-statistic calculated at each voxel and indicating the relationship between MNE amplitude estimations and the noise covariance (Dale et al. 2000). Finally, individual cortical surfaces were morphed onto a common brain, with 10,242 dipoles per hemisphere (Fischl et al. 1999), in order to compensate for inter-subject differences.

### Behavioral Data Analysis

The behavioral score was calculated as the average correct response across trials (4 multiple-choice question per narrative) for each participant. In order to verify that participants understood and completed the task, we performed a t-test between accuracy rates compared to chance-level (i.e., 0.33).Then, to test whether performance was affected by the type of task-irrelevant speech presented, we performed a paired t-test between the accuracy rates in both conditions. We additionally performed a median-split analysis of the behavioral scores across participant, based on their neural response to task-irrelevant speech (specifically the phrase-level response; see MEG data analysis), to test for possible interactions between performance on the to-be-attended speech and linguistic neural representation of task-irrelevant speech.

### MEG Data Analysis

#### Spectral Analysis

##### Scalp-level

Inter-trial phase coherence (ITPC) was calculated on the clean and segmented data. To this end, we applied a Fast Fourier Transform (FFT) to individual 8-second long segments and extracted the phase component at each frequency (from 0.1 to 15 Hz, with a 0.125Hz step). The normalized (z-scored) ITPC at each sensor was calculated using the Matlab circ_rtest function (circular statistics toolbox; Berens 2009). This was performed separately for the Structured and Non-Structured conditions. In order to determine which frequencies had significant ITPC, we performed a t-test between each frequency bin relative to the surrounding frequencies (average 2 bins from each side), separately for each condition (Nozaradan et al. 2018). In addition, we directly compared the ITPC spectra between the two conditions using a permutation test. In each permutation, the labels of the two conditions were randomly switched in half of the participants, and a paired t-test was performed. This was repeated 1,000 times, creating a null-distribution of t-values for this paired comparison, separately for each of the frequencies of interest. The t-values of the real comparisons were evaluated relative to this null-distribution, and were considered significant if they fell within the top 5% (one-way comparison, given our a-priori prediction that peaks in the Structured condition would be higher than in the Non-Structured condition). This procedure was performed on the average ITPC across all sensors, to avoid the need to correct for multiple comparisons, and focused specifically on four frequencies of interest (FOI): 4Hz, 2Hz, 1Hz and 0.5Hz which correspond to the four linguistic levels present in the Structured stimuli (syllables, words, phrases and sentences, respectively).

##### Source-level

Spectral analysis of source-level data was similar to that performed at the sensor level. ITPC was calculated for each frequency between 0.1 to 15 Hz (0.125Hz step) and a t-test between the ITPC at each frequency relative to the surrounding frequencies (2 bins from each side) was performed in order to validate the response peaks at the FOIs. Then, statistical comparison of responses at the source-level in the Structured and Non-Structured conditions focused only on the peaks that showed a significant difference between conditions at the scalp-level (in this case, the peak at 1Hz). In order to determine which brain-regions contributed to this effect we used 22 pre-defined ROIs in each hemisphere, identified based on multi-modal anatomical and functional parcellation (Glasser et al. 2016; Supplementary Neuroanatomical Results (table 1, page 180) and see Figure 3c). We calculated the mean ITPC value in each ROI, across participants and conditions, and tested for significant differences between them using a permutation test, which also corrected for multiple comparisons. As in the scalp-data, the permutation test was based on randomly switching the labels between conditions for half of the participants and conducting a paired t-test within each ROI. In each permutation we identified ROIs that passed a statistical threshold for a paired t-test (p<0.05) and the sum of their t-values was used as a global statistic. This procedure was repeated 1,000 times, creating a null-distribution for this global statistic. A similar procedure was applied to the real data, and if the global statistic (sum of t-values in the ROIs that passed an uncorrected threshold of p<0.05) fell within the top 5% of the null-distribution, the entire pattern could be considered statistically significant. This procedure was conducted separately within each hemisphere.

#### Speech-Tracking Analysis

Speech-tracking analysis was performed in order to estimate the neural response to the natural speech that served as the to-be-attended stimulus. To this end, we estimated the Temporal Response Function (TRF), which is a linear transfer-function expressing the relationship between features of the presented speech stimulus *s*(*t*) and the recorded neural response *r*(t). TRFs were estimated using normalized reverse correlation as implemented in the STRFpak Matlab toolbox (strfpak.berkeley.edu) and adapted for MEG data (Zion Golumbic, Cogan, et al. 2013). Tolerance and sparseness factors were determined using a jackknife cross-validation procedure, to minimize effects of over-fitting. In this procedure, given a total of *N* trials, a TRF is estimated between *s*(*t*) and *r*(*t*) derived from N-1 trials, and this estimate is used to predict the neural response to the left-out stimulus. The tolerance and sparseness factors that best predicted the actual recorded neural signal (predictive power estimated using Pearson’s correlation) were selected based on scalp-level TRF analysis (collapsed across conditions), and these were also used when repeating the analysis on the source-level data (David et al. 2007). The predictive power of the TRF model was also evaluated statistically by comparing it to a null-distribution obtain from repeating the procedure on 1,000 permutations of mismatched *s\**(*t*) and *r\**(*t*).

TRFs to the to-be-attended natural speech were estimated separately for trials in which the task-irrelevant speech was Structured vs. Non-Structured. TRFs in these two conditions were then compared statistically to evaluate the effect of the distractor type on neural encoding of to-be-attended speech. For scalp-level TRFs we used a spatial-temporal clustering permutation test to identify the time-windows where the TRFs differed significantly between conditions (fieldtrip; first level stat p<0.05, cluster corrected). We then turned to the source-level TRFs to further test which brain regions showed significant difference between conditions, by estimating TRFs in the same 22 pre-defined ROIs in each hemisphere used above. TRFs in the Structured vs. Non-Structured conditions were compared using t-tests in 20-ms long windows (focusing only on the 70-180ms time window which was found to be significant in the scalp-level analysis) and corrected for multiple comparisons using spatial-temporal clustering permutation test.

## Results

Data from one participant was excluded from all analyses due to technical problems during MEG recording. Six additional participants were removed only from source estimation analysis due to technical issues. The full behavioral and neural data are available at: https://osf.io/e93qa.

### Behavioral results

Behavioral results reflecting participants response accuracy on comprehension questions about narratives were significantly above chance [M=0.715, SD=±0.15; t(28)=26.67, p<0.001]. There were no significant differences in behavior as a function of whether task-irrelevant speech was Structured or Non-Structured [t(28)=-0.31, p=0.75; Figure 2). Additionally, to test for possible interactions between answering questions about the to-be-attended speech and linguistic neural representation of task-irrelevant speech, we performed a median-split analysis of the behavioral scores across participants. Specifically, we used the magnitude of the ITPC value at 1Hz in the Structured condition (averaged across all sensors), in order to split the sample into two groups – with high and low 1Hz responses. We performed a between-group t-test on the behavioral results in the Structured condition, and also on the difference between conditions (Structured – Non-Structured). Neither test showed significant differences in performance between participants whose 1Hz ITPC was above vs. below the median [Structured condition: t(27) = -1.07, p=0.29; Structured – Non-Structured: t(27) = -1.04, p=0.15]. Similar null-results were obtained when the median-split was based on the source-level data.

**Figure 2.**
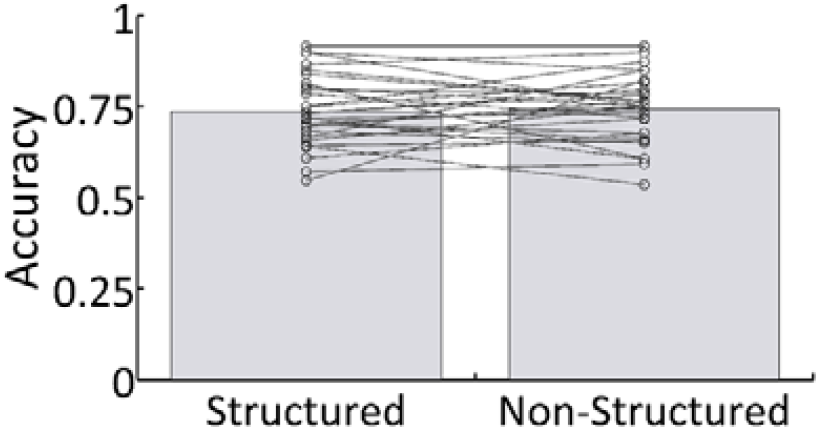
Behavioral results. Mean accuracy across all participants for both Structured and Non-Structured conditions. Lines represent individual results.

**Figure 3.**
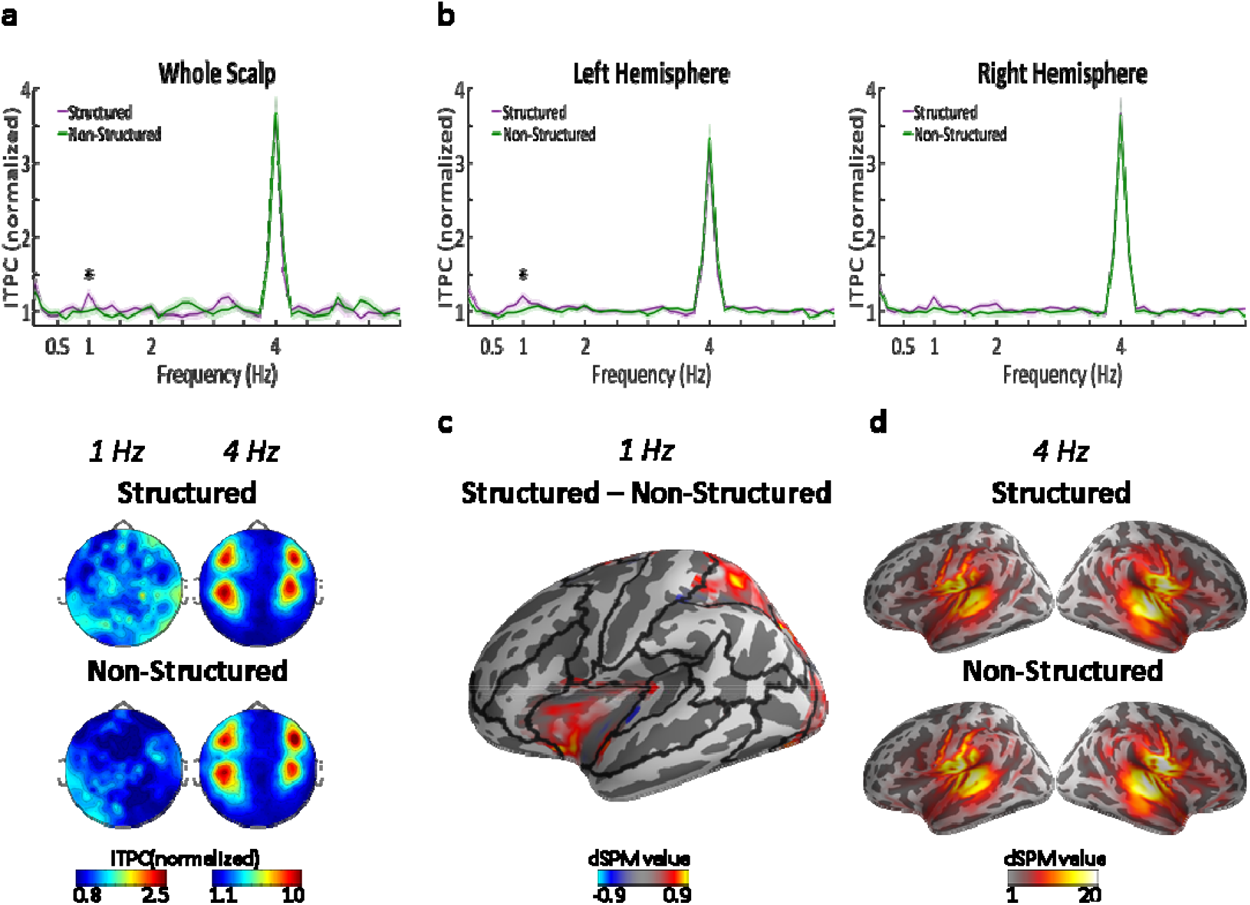
Neural tracking of linguistic structures in task-irrelevant speech. **a**, Top panel shows the ITPC spectrum at the scalp-level (average across all electrodes) in response to Structured (purple) and Non-Structured (green) task-irrelevant speech. ITPC values z-score normalized, as implemented in the circ_rtest function (see Methods). Shaded areas indicate SEM across participants (n=29). Asterisk represents statistically significant difference (p<0.05) between conditions, indicating a significant response at 1Hz for Structured task-irrelevant speech, which corresponds to the phrase-level. Bottom panel shows the scalp-topography of ITPC at 4Hz and 1Hz in the two conditions. **b**, ITPC spectrum at the source-level, averaged across all voxels in each hemisphere. Shaded highlights denote SEM across participants (n=23). Asterisk represents statistically significant difference (p<0.05) between conditions, indicating a significant response at 1Hz for Structured task-irrelevant speech in the left, but not right hemisphere. **c**, Source-level map on a central inflated brain depicting the ROIs in the left hemisphere where significant differences in ITPC at 1Hz were found for Structured vs. Non-Structured task-irrelevant speech. Black lines indicate the parcellation into the 22 ROIs used for source-level analysis. **d**, Source-level maps showing localization of the syllabic-rate response (4Hz) in both conditions.

### Hierarchical frequency-tagged responses to task-irrelevant speech

Scalp-level spectra of the Inter-trial phase coherence (ITPC) showed a significant peak at the syllabic-rate (4Hz) in response to both Structured and Non-Structured hierarchical frequency-tagged speech, with a 4-pole scalp-distribution common to MEG recorded auditory responses (Figure 3a) (p< 10^-9; large effect size, Cohen’s d > 1.5 in both). As expected, there was no significant difference between Structured and Non-Structured condition in the 4Hz response (p=0.899). Importantly, we also observed a significant peak at 1Hz in the Structured condition (p<0.003; moderate effect size, Cohen’s d = 0.6), but not in the Non-Structured condition (p=0.88). Comparison of the 1Hz ITPC between these conditions also confirmed a significant difference between them (p=0.045; moderate effect size, Cohen’s d = 0.57). The scalp-distribution of the 1Hz peak did not conform to the typical auditory response topography, suggesting different neural generators. No other significant peaks were observed at any other frequencies, including the 2Hz or 0.5Hz word and sentence-level rates, nor did the responses at these rates differ significantly between conditions.

In order to determine the neural source of the 1Hz peak in the Structured condition, we repeated the spectral analysis in source-space. An inverse solution was applied to individual trials and the ITPC was calculated in each voxel. As shown in Figure 3b, the source-level ITPC spectra, averaged over each hemisphere separately, is qualitatively similar to that observed at the scalp-level. The only significant peaks were at 4Hz in both conditions (p< 10^−8^; large effect size, Cohen’s d > 1.7 in both conditions and both hemispheres) and at 1Hz in the Structured condition (left hemisphere p=0.052, Cohen’s d = 0.43; right hemisphere p<0.007, Cohen’s d = 0.6), but not in the Non-Structured condition. Statistical comparison of the 1Hz peak between conditions revealed a significant difference between the Structured and Non-Structured condition over the left hemisphere (p=0.026, Cohen’s d = 0.57), but not over the right hemisphere (p=0.278).

Figure 3c shows the source-level distribution within the left hemisphere of the difference in 1Hz ITPC between the Structured and Non-Structured condition. The effect was observed primarily in frontal and parietal regions. Statistical testing evaluating the difference between conditions was performed in 22 pre-determined ROIs per hemisphere, using a permutation test. This indicated significant effects in several ROIs in the left hemisphere including the inferior-frontal cortex and superior parietal cortex, as well as the mid-cingulate and portions of the middle and superior occipital gyrus (cluster-corrected p=0.002). No ROIs survived multiple-comparison correction in the right hemisphere (cluster-corrected p=0.132), although some ROIs in the right cingulate were significant at an uncorrected level (p<0.05).

With regard to the 4Hz peak, it was localized as expected to bilateral auditory cortex and did not differ significantly across conditions in either hemisphere (left hemisphere: p=0.155, right hemisphere: p=0.346). We therefore did not conduct a more fine-grained analysis of different ROIs. As in the scalp-level data, no peaks were observed at 2Hz and no significant difference between conditions (left hemisphere: p=0.963, right hemisphere: p=0.755).

### Speech Tracking of to-be-attended speech

Speech tracking analysis of responses to the to-be-attended narrative yielded robust TRFs (scalp-level predictive power r=0.1, p<0.01 vs. permutations). The TRF time-course featured two main peaks, one ∼80ms and the other ∼140ms, in line with previous TRF estimations (Akram et al. 2017; Fiedler et al. 2019; Brodbeck et al. 2020b). Both the scalp-level and source-level TRF analysis indicated that TRFs were predominantly auditory – showing the common 4-pole distribution at the scalp-level (Figure 4a) and in the source-level analysis was localized primarily to auditory cortex (superior temporal gyrus and sulcus; STG/STS) as well as left insula/IFG (Figure 4b; 140ms).

**Figure 4.**
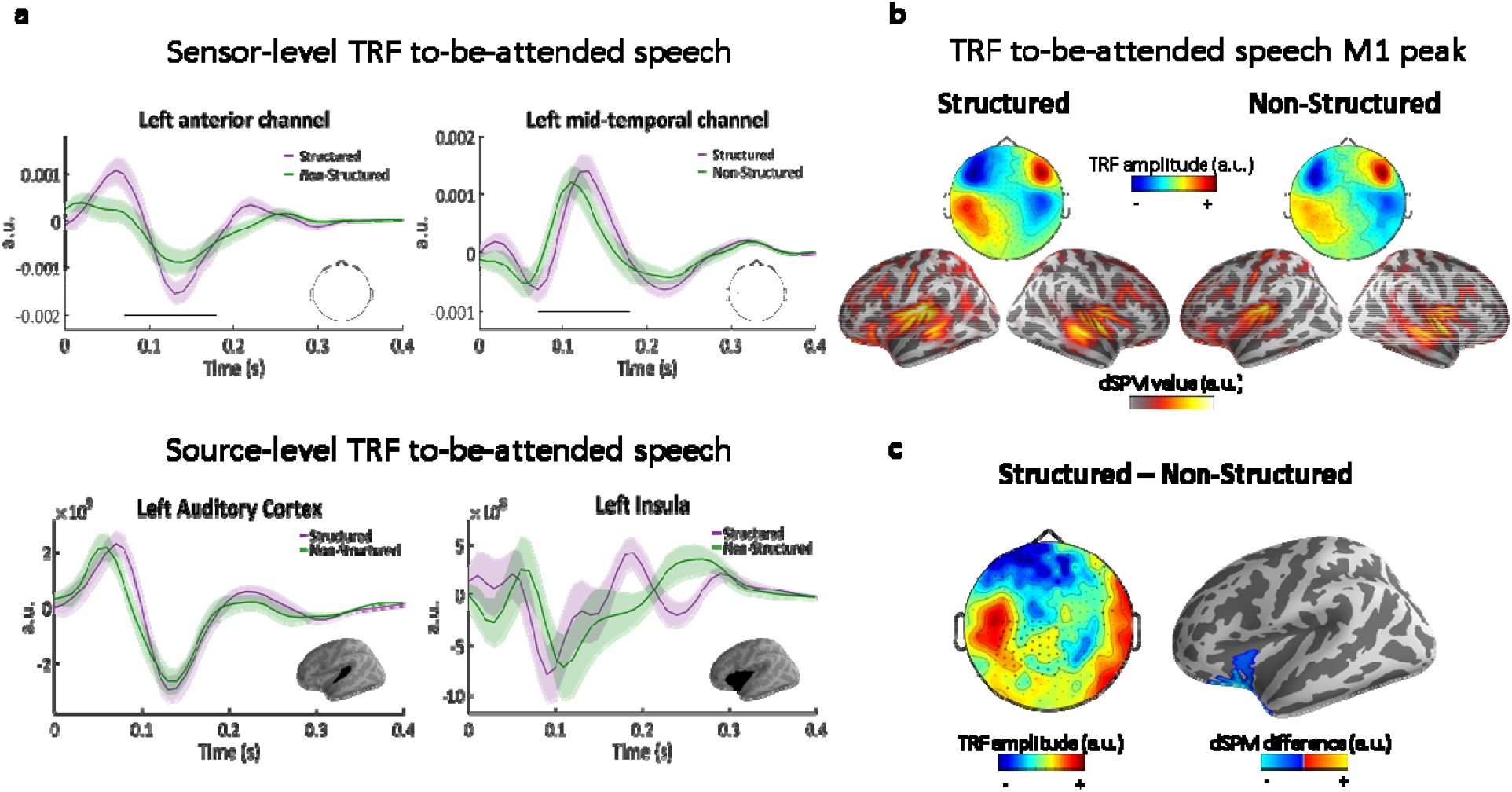
**a, top**, TRF examples from two sensors, showing the positive and negative poles of the TRF over the left hemisphere. Shaded highlights denote SEM across participants. Black line indicates the time points where the difference between conditions was significant (spatio-temporal cluster corrected). **bottom**, TRF examples from source-level ROIs in left Auditory Cortex and in the left Inferior Frontal/Insula region. **b**, topographies (top) and source estimations (bottom) for each condition at M1 peak (140 ms). **c, left**, topography of the difference between conditions (Structured – Non-Structured) at the M1 peak (140 ms). Asterisks indicate the MEG channels where this difference was significant (cluster corrected). **right**, significant cluster at the source-level.

When comparing the TRFs to the to-be-attended speech as a function of whether the competing task-irrelevant stimulus was Structured vs. Non-Structured, some interesting differences emerged. Spatial-temporal clustering permutation test on the scalp-level TRFs revealed significant differences between the conditions between 70-180 ms (p<0.05; cluster corrected), including both the early and late TRF peaks, at a large number of sensors primarily on the left (Figure 4a&c). Specifically, TRF responses to the to-be-attended speech were enhanced when the task-irrelevant stimulus was Structured condition vs. Non-Structured. The effect was observed at the scalp-level with opposite polarity in frontal vs. medial sensors, and was localized at the source-level to a single cluster in the left inferior-frontal cortex, that included portions of the insula and orbito-frontal cortex (Figure 4c; spatial-temporal clustering p=0.0058).

## Discussion

In this MEG study, frequency-tagged hierarchical speech was used to probe the degree to which linguistic processing is applied to task-irrelevant speech, and how this interacts with processing speech that is presumably in the focus of attention (‘to-be-attended’). As expected, we observe obligatory acoustic representation of task-irrelevant speech, regardless of whether it was Structured or Non-Structured, which manifest as a 4Hz syllable-level response localized to bilateral auditory regions in the STG/STS. Critically, for Structured task-irrelevant speech we also find evidence for neural tracking of the phrasal structure, with a 1Hz peak localized primarily to left inferior-frontal cortex and left posterior-parietal cortex. The regions are not associated with low-level processing, but rather play an important role in speech processing and higher order executive functions (Dronkers et al. 2004; Humphries et al. 2006; Linden 2007; Corbetta et al. 2008; Edin et al. 2009; Hahn et al. 2018). Additionally, we find that the speech-tracking response to the *to-be-attended* speech in left inferior frontal cortex was also affected by whether task-irrelevant speech was linguistically Structured or not. These results contribute to ongoing debates regarding the nature of the competition for processing resources during speech-on-speech masking, demonstrating that linguistic processes requiring integration of input over relatively long timescales can indeed be applied to task-irrelevant speech.

### The debate surrounding linguistic processing of task-irrelevant speech

Top-down attention is an extremely effective process, by which the perceptual and neural representation of task-relevant speech are enhanced at the expense of task-irrelevant stimuli, and speech in particular (Horton et al. 2013; Zion Golumbic, Ding, et al. 2013; O’Sullivan et al. 2015; Fiedler et al. 2019; Teoh and Lalor 2019). However, the question still stands: what degree of linguistic processing is applied to task-irrelevant speech? One prominent position is that attention is required for linguistic processing and therefore speech that is outside the focus of attention is not processed beyond its sensory attributes (Lachter et al. 2004; Brodbeck, Hong, et al. 2018; Ding et al. 2018). However, several lines of evidence suggest that linguistic features of task-irrelevant speech can be processed as well, at least under certain circumstances. For example, task-irrelevant speech is more disruptive to task performance if it is intelligible, as compared to unintelligible noise-vocoded or rotated speech (Marrone et al. 2008; Iyer et al. 2010; Best et al. 2012; Gallun and Diedesch 2013; Swaminathan et al. 2015; Kidd et al. 2016) or a foreign language (Freyman et al. 2001; Rhebergen et al. 2005; Cooke et al. 2008; Calandruccio et al. 2010; Francart et al. 2010). This effect, referred to as informational masking, is often attributed to the detection of familiar acoustic-phonetic features in task-irrelevant speech, that can lead to competition for phonological processing (‘phonological interference’) (Durlach et al. 2003; Drullman and Bronkhorst 2004; Kidd et al. 2008; Shinn-Cunningham 2008; Rosen et al. 2013). However, the phenomenon of informational masking alone is insufficient for determining the extent to which task-irrelevant speech is processed beyond identification of phonological units.

Other lines of investigation have focused more directly on whether task-irrelevant speech is represented at the semantic level. Findings that individual words from a task-irrelevant source are occasionally detected and recalled, such as one’s own name, (Cherry 1953; Wood and Cowan 1995; Conway et al. 2001; Röer et al. 2017b, 2017a), have been taken as evidence that task-irrelevant inputs can be semantically processed, albeit the information may not be consciously available. Along similar lines, a wealth of studies demonstrate the ‘Irrelevant Sound Effect’ (ISE), showing that the semantic content of task-irrelevant input affects performance on a main task, mainly through priming effects and interference with short-term memory (Lewis 1970; Bentin et al. 1995; Surprenant et al. 1999; Dupoux et al. 2003; Beaman 2004; Rivenez et al. 2006; Beaman et al. 2007; Rämä et al. 2012; Aydelott et al. 2015; Schepman et al. 2016; Vachon et al. 2019). However, an important caveat precludes interpreting these findings as clear-cut evidence for semantic processing of task-irrelevant speech: Since these studies primarily involve presentation of arbitrary lists of words (mostly nouns), usually at a relatively slow rate, an alternative explanation is that the ISE is simply a result of occasional shifts of attention towards task-irrelevant stimuli (Carlyon 2004; Lachter et al. 2004). Similarly, the effects of informational masking discussed above can also be attributed to a similar notion of perceptual glimpsing, i.e., gleaning bits of the task-irrelevant speech in the short ‘gaps’ in the speech that is to-be-attended (Cooke 2006; Kidd et al. 2016; Fogerty et al. 2018). These claims – that effects of task-irrelevant speech are not due to parallel processing but reflect shifts of attention – are extremely difficult to reject empirically, as they would require insight into the listeners’ internal state of attention, which at present is not easy to operationalize.

### Phrase-level response to task-irrelevant speech

In attempt to broach the larger question of processing task-irrelevant speech, the current study takes a different approach by focusing not on detection of single words, but on linguistic processes that operate over longer timescales. To this end the stimuli used here, in both the to-be-attended and the task-irrelevant ear, was continuous speech rather than word-lists whose processing requires accumulating and integrating information over time, which is strikingly different than the point-process nature of listening to word-lists (Fedorenko et al. 2016). Using continuous speech is also more representative of the type of stimuli encountered naturally in the real world (Hill and Miller 2010; Risko et al. 2016; Matusz et al. 2019; Shavit-Cohen and Zion Golumbic 2019). In addition, by employing hierarchical frequency-tagging, we were able to obtain objective and direct indications of which levels of information were detected within task-irrelevant speech. Indeed, using this approach we were able to identify a phrase-level response for Structured task-irrelevant speech, which serves as a positive indication that these stimuli are indeed processed in a manner sufficient for identifying the boundaries of syntactic structures.

An important question to ask is whether the phrase-level response observed for task-irrelevant speech can be explained by attention shifts? Admittedly, in the current design participants could shift their attention between streams in an uncontrolled fashion, allowing them to ‘glimpse’ portions of the task-irrelevant speech, integrate and comprehend (portions of) it. Indeed, this is one of the reasons we refrain from referring to the task-irrelevant stream as “unattended”: since we have no principled way to empirically observe the internal loci or spread of attention, we chose to focus on its behavioral relevance rather than make assumptions regarding the participants’ attentional-state. Despite the inherent ambiguity regarding the underlying dynamics of attention, the fact that here we observe a phrase-level response for task-irrelevant speech is direct indication that phonetic-acoustic information from this stream was decoded and integrated over time, allowing the formation of long-scale representations for phrasal boundaries. If this is a result of internal ‘glimpsing’, this would imply either that (a) the underlying hierarchical speech structure was detected and used to guide ‘glimpses’ in a rhythmic-manner to points in time that are most informative; or (b) that ‘glimpses’ occur irregularly, but that sufficient information is gleaned through them and stored in working-memory to allow the consistent detection phrasal boundaries in task-irrelevant speech. Both of these options imply a sophisticated multiplexed encoding-scheme for successful processing of concurrent speech, that relies on precise temporal control and working-memory storage. Another possibility, of course, is that there is no need for attention-shifts and that the system has sufficient capacity to process task-irrelevant speech in parallel to focusing primarily on the to-be-attended stream. As mentioned above, the current data cannot provide insight into which of these listening-strategies underlies the generation of the observed phrase-level response to task-irrelevant speech. However, we hope that future studies will gain empirical access into the dynamic of listeners’ internal attentional state and help shed light on this pivotal issue.

The current study is similar in design to another recent study by Ding et al. (2018) where Structured frequency-tagged speech was presented as a task-irrelevant stimulus. In contrast to the results reported here, they did not find significant peaks at any linguistic-related frequencies in the neural response to task-irrelevant speech. In attempt to resolve this discrepancy, it is important to note that these two studies differ in an important way – in the listening effort that was required of participants in order to understand the to-be-attended speech. While in the current experiment to-be-attended speech was presented in its natural form, mimicking the listening effort of real-life speech-processing, in the study by Ding et al. (2018) to-be-attended speech was time-compressed by a factor of 2.5 and naturally occurring gaps were removed, making the comprehension task substantially more effortful (Nourski et al. 2009; Müller et al. 2019). Load Theory of Attention proposes that the allocation of processing resources among competing inputs can vary as a function of the perceptual traits and cognitive load imposed by the task (Lavie et al. 2004; Murphy et al. 2017). Accordingly, it is plausible that these divergent results are due to the extreme difference in the perceptual load and listening effort in the two studies. Specifically, if understanding the to-be-attended speech imposes relatively low perceptual and cognitive load, then sufficient resources may be available to additionally process aspects of task-irrelevant speech, but that this might not be the case as the task becomes more difficult and perceptually demanding (Wild et al. 2012; Gagné et al. 2017; Peelle 2018).

More broadly, the comparison between these two studies invites re-framing of the question regarding the type / level of linguistic processing applied to task-irrelevant speech, and propels us to think about this issue not as a yes-or-no dichotomy, but perhaps as a more flexible process that depends on the specific context (Brodbeck et al. 2020b). The current results provide a non-trivial positive example for processing task-irrelevant speech that is indeed processed beyond its acoustic attributes, in an experimental context that closely emulates the perceptual and cognitive load encountered in real-life (despite the admitted unnatural nature of the task-irrelevant speech). At the same time, they do not imply that this is always the case, as is evident from the diverse results reported in the literature regarding processing task-irrelevant speech, as discussed at length above. Rather, they invite adopting a more flexible perspective of processing bottlenecks within the speech processing system, that takes into consideration the perceptual and cognitive load imposed in a given context, in line with load theory of attention (Mattys et al. 2012; Lavie et al. 2014; Fairnie et al. 2016; Gagné et al. 2017; Peelle 2018). Supporting this perspective, others have also observed that the level of processing applied to task-irrelevant stimuli can be affected by task demands (Hohlfeld and Sommer 2005; Pulvermüller et al. 2008). Moreover, individual differences in attentional abilities, and particularly the ability to process concurrent speech, have been attributed partially to working-memory capacity, a trait associated with the availability of more cognitive resources (Beaman et al. 2007; Forster and Lavie 2008; Naveh-Benjamin et al. 2014; Lambez et al. 2020) but cf. (Elliott and Briganti 2012). As cognitive neuroscience research increasingly moves towards studying speech processing and attention in real-life circumstances, a critical challenge will be to systematically map out the perceptual and cognitive factors that contribute to, or hinder, the ability to glean meaningful information from stimuli that are outside the primary focus of attention.

### The brain regions where phrase-level response is observed

The phrase-level neural response to task-irrelevant Structured speech was localized primarily to two left-lateralized clusters: one in the left anterior fronto-temporal cortex and the other in left posterior-parietal cortex. The fronto-temporal cluster, which included the IFG and insula, is known to play an important role in speech processing (Dronkers et al. 2004; Humphries et al. 2006; Brodbeck, Presacco, et al. 2018; Blank and Fedorenko 2020). The left IFG and insula are particularly associated with linguistic processes that require integration over longer periods of time, such as syntactic structure building and semantic integration of meaning (Fedorenko et al. 2016; Matchin et al. 2017; Schell et al. 2017), and are also recruited when speech comprehension requires effort, such as for degraded or noise-vocoded speech (Davis and Johnsrude 2003; Obleser and Kotz 2010; Davis et al. 2011; Hervais-Adelman et al. 2012). Accordingly, observing a phrase-level response to task-irrelevant speech in these regions is in line with their functional involvement in processing speech under adverse conditions.

With regard to the left posterior-parietal cluster, the interpretation for why a phrase-level response is observed there is less straightforward. Although some portions of the parietal cortex are involved in speech processing, these are typically more inferior than the cluster found here (Hickok and Poeppel 2007; Smirnov et al. 2014). However, both the posterior-parietal cortex and inferior frontal gyrus play an important role in verbal working-memory (Todd and Marois 2004; Postle et al. 2006; Linden 2007; McNab and Klingberg 2008; Edin et al. 2009; Østby et al. 2011; Rottschy et al. 2012; Gazzaley and Nobre 2012; Ma et al. 2012; Meyer et al. 2014, 2015; Yue et al. 2019; Fedorenko and Blank 2020). Detecting the phrasal structure of task-irrelevant speech, while focusing primarily on processing the to-be-attended narratives, likely requires substantial working-memory for integrating chunks of information over time. Indeed, attention and working-memory are tightly linked constructs (McNab and Klingberg 2008; Gazzaley and Nobre 2012; Vandierendonck 2014), and as mentioned above, the ability to control and maintain attention is often associated with individual working-memory capacity (Cowan et al. 2005; Beaman et al. 2007; Forster and Lavie 2008; Naveh-Benjamin et al. 2014; Lambez et al. 2020). Therefore, one possible interpretation for the presence of a phrase-level response to task-irrelevant speech in the left posterior-parietal cortex and inferior frontal regions, is their role in forming and maintaining a representation of task-irrelevant stimuli in working-memory, perhaps as a means for monitoring the environment for potentially important events.

### Why no word-level response?

Although in the current study we found significant neural response to task-irrelevant speech at the phrasal-rate, we did not see peaks at the word- or at the sentence-rate. Regarding the sentence-level response, it is difficult to determine whether the lack of an observable peak indicates that the stimuli were not parsed into sentences, or if this null-result is due to the technical difficulty of obtaining reliable peaks at low-frequencies (0.5Hz) given the 1/f noise-structure of neurophysiological recordings (Pritchard 1992; Miller et al. 2009). Hence, this remains an open question for future studies. Regarding the lack of a word-level response at 2Hz for Structured task-irrelevant stimuli, this was indeed surprising, since in previous studies using the same stimuli in a single-speaker context we observe a prominent peak at *both* the word- and the phrase-rate (Makov et al. 2017). Although we do not know for sure why the 2Hz peak is not observed when this speech was presented as task-irrelevant concurrently with another narrative, we can offer some speculations for this null-result: One possibility is that the task-irrelevant speech was indeed parsed into words as well, but that the neural signature of 2Hz parsing was not observable due to interference from the acoustic contributions at 2Hz (see Supplementary Materials and (Luo and Ding 2020). However, another possibility is that the lack of a word-level response for task-irrelevant speech indicates that it does not undergo full lexical analysis. Counter to the linear intuition that syntactic structuring depends on identifying individual lexemes, there is substantial evidence that lexical and syntactic processes are separable and dissociable cognitive processes, that rely on partially different neural substrates (Friederici and Kotz 2003; Hagoort 2003; Humphries et al. 2006; Nelson et al. 2017; Schell et al. 2017; Pylkkänen 2019; Morgan et al. 2020). Indeed, a recent frequency-tagging study showed that syntactic phrasal structure can be identified (generating a phrase-level peak in the neural spectrum) even in the complete absence of lexical information (Getz et al. 2018). Hence, it is possible that when speech is task-irrelevant and does not receive full attention, it is processed only partially, and that although phrasal boundaries are consistently detected, task-irrelevant speech does not undergo full lexical analysis. This matter regarding the depth of lexical processing of task-irrelevant speech, and its interaction with syntactic analysis, remains to be further explored in future research.

### Task-irrelevant influence on processing to-be-attended speech

Besides analyzing the frequency-tagged neural signatures associated with encoding the ***task-irrelevant stimuli***, we also looked at how the neural encoding of ***to-be-attended speech*** was affected by the type of task-irrelevant speech it was paired with. In line with previous MEG studies, the speech-tracking response (estimated using TRFs) was localized to auditory temporal regions bilaterally and left inferior frontal regions (Ding and Simon 2012; Zion Golumbic, Cogan, et al. 2013; Puvvada and Simon 2017). The speech tracking response in auditory regions was similar in both conditions, however the response in left inferior-frontal cortex was modulated by the type of task-irrelevant speech presented and was enhanced when task-irrelevant speech was Structured vs. when it was Non-Structured. This pattern highlights the nature of the competition for resources triggered by concurrent stimuli. When the task-irrelevant stimulus was Non-Structured, even though it was comprised of individual phonetic-acoustic units, it did not contain meaningful linguistic information and therefore did not require syntactic and semantic resources. However, the Structured task-irrelevant speech poses more of a competition, since it constitutes fully intelligible speech. Indeed, it is well established that intelligible task-irrelevant speech causes more competition and therefore are more distracting than non-intelligible speech (Rhebergen et al. 2005; Iyer et al. 2010; Best et al. 2012; Gallun and Diedesch 2013; Carey et al. 2014; Kilman et al. 2014; Swaminathan et al. 2015; Kidd et al. 2016). A recent EEG study found that responses to both target and distractor speech are enhanced when the distractor was intelligible vs. unintelligible (Olguin et al. 2018), although this may depend on the specific type of stimulus used (Rimmele et al. 2015). However, in most studies it is difficult to ascertain the level(s) of processing where competing between the inputs occurs, and many effects can be explained by variation in the acoustic nature of maskers (Ding and Simon 2014). The current study is unique in that all low-level features of Structured and Non-Structured speech stimuli were perfectly controlled, allowing us to demonstrate that interference goes beyond the phonetic-acoustic level and also occurs at higher linguistic levels. The findings that the speech tracking response of the to-be-attended narratives is enhanced when competing with a Structured task-irrelevant speech, specifically left inferior-frontal brain regions, where we also observed tracking of the phrase-structure of task-irrelevant speech, pinpoints the locus of this competition to these dedicated speech-processing regions, above and beyond any sensory-level competition (Davis et al. 2011; Brouwer et al. 2012; Hervais-Adelman et al. 2012). Specifically, they suggests that the enhanced speech tracking response in IFG reflects the investment of additional listening effort for comprehending the task-relevant speech (Vandenberghe et al. 2002; Gagné et al. 2017; Peelle 2018).

Since the neural response to the to-be-attended speech was modulated by the type of competition it faced, then why was this not mirrored in the current behavioral results as well? In the current study participants achieved similar accuracy rates on the comprehension questions regardless of whether the natural-narratives were paired with Structured or Non-Structured stimuli in the task-irrelevant ear, and there was no significant correlation between the neural effects and performance. We attribute the lack of a behavioral effect primarily to the insensitivity of the behavioral measures used here, that consisted of asking four multiple-choice questions after each 45-second long narrative. Although numerous previous studies have been able to demonstrate behavioral ‘intrusions’ of the task-irrelevant stimuli on performance of an attended-task, these have been shown using more constrained experimental paradigms, that have the advantage of probing behavior at a finer scale, but are substantially less ecological (e.g., memory-recall for short lists of words or priming effects; (Tun et al. 2002; Dupoux et al. 2003; Rivenez et al. 2006, 2008; Carey et al. 2014; Aydelott et al. 2015). In moving towards studying speech processing and attention under more ecological circumstances, using natural continuous speech, we face an experimental challenge of obtaining sufficiently sensitive behavior measures without disrupting listening with an ongoing task (e.g., target detection) or encroaching too much on working-memory. This is a challenge shared by many previous studies similar to ours, and is one of the main motivations for turning directly to the brain and studying neural activity during uninterrupted listening to continuous speech, rather than relying on sparse behavioral indications (Ding et al. 2016; Makov et al. 2017; Brodbeck, Hong, et al. 2018; Broderick et al. 2018, 2019; Donhauser and Baillet 2019).

## Conclusions

The current study contributes to ongoing efforts to understand how the brain deals with the abundance of auditory inputs in our environment. Our results indicate that even though top-down attention effectively enables listeners to focus on a particular task-relevant source of input (speech in this case), this prioritization can be affected by the nature of task-irrelevant sounds. Specifically, we find that when the latter constitutes meaningful speech, left fronto-temporal speech-processing regions are engaged in processing both stimuli, potentially leading to competition for resources and more effortful listening. Additional brain regions, such as the PPC, are also engaged in representing some aspects of the linguistic structure of task-irrelevant speech, which we interpret as maintaining a representation of what goes on in the ‘rest of the environment’, in case something important arises. Importantly, similar interactions between the structure of task-irrelevant sounds and responses to the to-be-attended sounds have been previously demonstrated for non-verbal stimuli as well (Makov and Zion Golumbic 2020). Together, this highlights the fact that attentional selection is not an all-or-none processes, but rather is a dynamic process of balancing the resources allocated to competing input, which is highly affected by the specific perceptual, cognitive and environmental aspects of a given task.

## Acknowledgements

This work was funded by Binational Science Foundation (BSF) grant # 2015385 and ISF grant #2339/20. We would like to thank Dr. Nai Ding for helpful comments on a previous version of this paper.

## Supplementary Materials

The modulation spectrum of Structured speech stimuli used in this study featured a prominent peak at 4Hz, corresponding to the syllable-rate, and an additional smaller peak at 2Hz. Since the stimuli were built by concatenating 250-ms long syllables, while taking care **not** to introduce any additional acoustic events that would introduce other rhythmic regularities (such as systematic gaps between words; Buiatti et al. 2009), we hypothesized that it may be related to the **order** of the syllables within the Structured sequences. Specifically, since our Structured stimuli was comprised of bi-syllabic Hebrew words, there may be a systematic difference in the envelope-shape of syllables at the beginning vs. end of words. For example, in the materials used here, it was indeed more common to start a word with a CV syllable than to end with one (Figure S1 and S2a). This, in turn, could lead to subtle yet systematic differences in the envelope-shape at even vs. odd positions in the stimulus, particularly after averaging across sentences/trials, resulting in a 2Hz peak in the modulation spectrum. A recent study by Luo and Ding (2020) nicely demonstrates that an acoustic-driven 2Hz peak can be induced simply by amplifying every second syllable in multi-syllable words.

**Figure S1.**
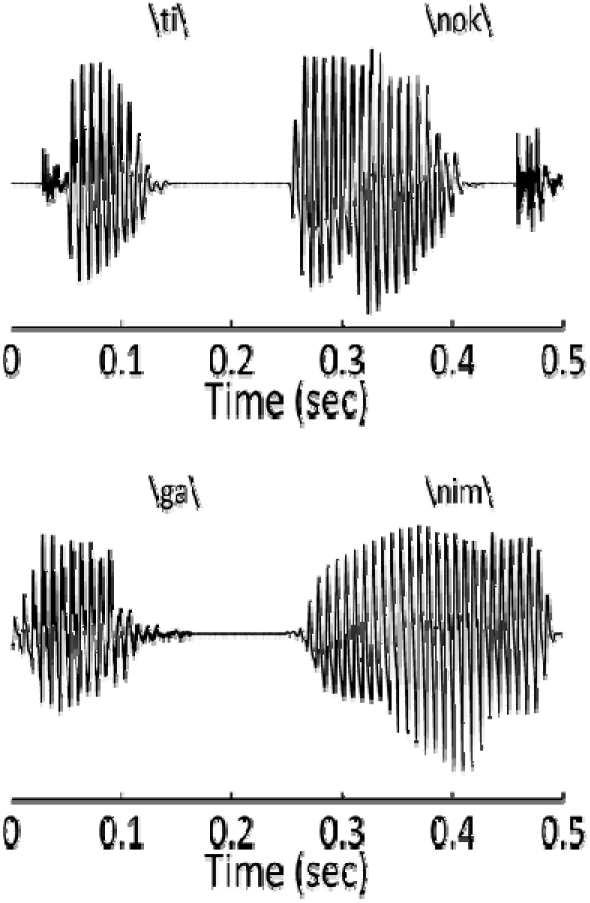
Two examples of start and end syllables for bi-syllabic Hebrew words. This illustrates the natural differences in envelope-shape for different syllables. If short CV syllables systematically occur at the start vs. end of words, this can potentially create a 2Hz peak in the modulation spectrum.

To better understand the origin of the 2Hz peak in our Structured stimuli we ran several simulations, testing how the order of the syllables within the sequence affected the modulation spectrum. First, we created Position-Controlled Stimuli (Figure S2b), which were pseudo-sentences comprised of the same syllables as the Structured speech, but ordered in a manner that did not create linguistic meaning. Importantly, randomization was performed in a manner so that each syllable maintained the same position it had in the original sentences. For example, if the syllable /gu/ was the first in original Sentence #1 and the syllable /buk/ was the last, then in the Position-Controlled stimuli these two syllables will still be first or last, respectively, but will no longer be part of the same pseudo-sentence (see concrete examples in Figure S2b). In this manner, the average audio-envelope across sentences is still identical to the Structured materials, but the stimuli no longer carry linguistic meaning.

The same procedure was used for calculating the modulation spectrum as were applied to the original Structured stimuli (see Methods for full details). Briefly, a total of 52 pseudo-sentences were constructed and randomly concatenated to form 50 sequences (56-seconds long). To stay faithful to the analysis procedure applied later to the MEG data, sequences were then divided into 8-second epochs, the envelope was extracted and FFT was applied to each epoch. The modulation spectrum is the result of averaging the spectrum across all epochs. We find that, indeed, the modulation spectrum of the original Structured materials and the Position-Controlled materials are basically identical, both containing similar peaks at 4Hz and 2Hz. This supports our hypothesis that the 2Hz peak stemmed from a natural asymmetry in the type of syllables that occur in start vs. end positions of bi-syllabic words (at least in Hebrew). We note that this type of Position-Controlled stimuli were used by us as Non-Structured stimuli in a previous study (Makov et al. 2017), and this is likely a more optimal choice as a control stimuli for future studies, as it allows to more confidently attribute differences at 2Hz in the neural response between Structured and Position-Controlled Non-Structured stimuli to linguistic, rather than acoustic, effects.

We next ran two additional simulations to determine what form of syllable randomization eliminates the 2Hz peak. We found that when creating pseudo-sentences where syllables are completely randomized and not constrained by position, the 2Hz peak is substantially reduced (Figure S2c). However, in this case we used a Fixed set of 52 pseudo-sentences to form sequences of stimuli. When we further relaxed this constraint, and allowed different randomization in different sequences (Non-Fixed Randomized stimuli; Figure S2d), the 2Hz peak was completely eliminated. The latter is akin to the Non-Structured stimuli used in the current experiment, and hence they were, in fact, not fully controlled at the acoustic level for 2Hz modulations.

That said, since we did not in fact see any peak in the neural data at 2Hz, and the effect we did find was at 1Hz (which *is* controlled across conditions), this caveat does not affect the validity of the results reported here. Future studies using this hierarchical frequency-tagging approach should take care to equate the modulation spectrum of experimental and control stimuli on this dimension as well (as was done by Makov et al. 2017).

**Figure S2.**
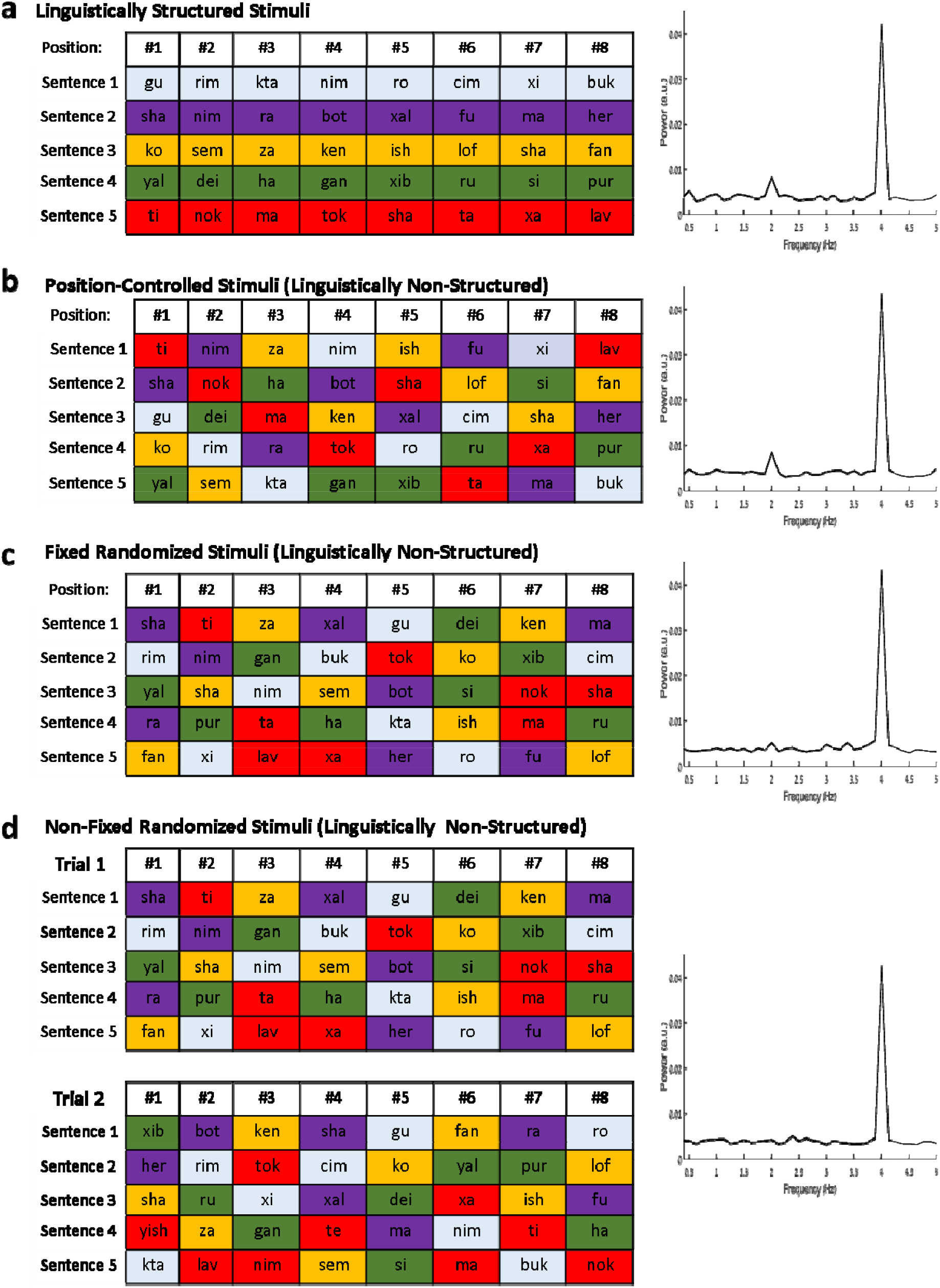
The effects of syllable-order on the modulation spectrum. Each panel shows examples for sentences/pseudo-sentence constructed in different ways (left) and the resulting modulation spectrum for this type of stimulus (right). a, Examples of five Structured sentences used in the current study. b, Example of Position-Controlled pseudo-sentences, where the syllables used in the Structured sentences are randomized across pseudo-sentences, but maintain the same position as in the Structured sentences. c, Examples of Fixed Randomized Stimuli, where a set of pseudo-sentences are constructed through random assignment of syllables (with no position-constraints), and these fixed pseudo-sentences are used to construct all the stimuli across trials. d, Examples of Non-Fixed Randomized Stimuli, where different randomization of syllables is used to construct stimuli in each trial.

## References

Akram S, Simon JZ, Babadi B. 2017. Dynamic estimation of the auditory temporal response function from MEG in competing-speaker environments. IEEE Trans Biomed Eng. 64:1896– 1905.

Ambler BA, Fisicaro SA, Proctor RW. 1976. Temporal characteristics of primary-secondary message interference in a dichotic listening task. Mem Cognit. 4:709–716.

Aydelott J, Jamaluddin Z, Nixon Pearce S. 2015. Semantic processing of unattended speech in dichotic listening. J Acoust Soc Am. 138:964–975.

Beaman CP. 2004. The Irrelevant Sound Phenomenon Revisited: What Role for Working Memory Capacity? J Exp Psychol Learn Mem Cogn. 30:1106–1118.

Beaman CP, Bridges AM, Scott SK. 2007. From Dichotic Listening to the Irrelevant Sound Effect: a Behavioural and Neuroimaging Analysis of the Processing of Unattended Speech. Cortex. 43:124–134.

Bentin S, Kutas M, Hillyard SA. 1995. Semantic processing and memory for attended and unattended words in dichotic listening: Behavioral and electrophysiological evidence. J Exp Psychol Hum Percept Perform. 21:54–67.

Berens P. 2009. CircStatil: A MATLAB Toolbox for Circular Statistics. J Stat Softw. 31:1–21.

Best V, Marrone N, Mason CR, Kidd G. 2012. The influence of non-spatial factors on measures of spatial release from masking. J Acoust Soc Am. 131:3103–3110.

Blank IA, Fedorenko E. 2020. No evidence for differences among language regions in their temporal receptive windows. Neuroimage. 219:116925.

Boersma P. 2011. Praatil: doing phonetics by computer [Computer program]. http://www.praat.org/.

Broadbent DE. 1958. Perception and Communication. London: Pergamon Press.

Brodbeck C, Hong LE, Simon JZ. 2018. Rapid Transformation from Auditory to Linguistic Representations of Continuous Speech. Curr Biol. 28:3976-3983.e5.

Brodbeck C, Jiao A, Hong LE, Simon JZ. 2020a. Dynamic Processing of Background Speech 1 at the Cocktail Party: Evidence for Early 2 Active Cortical Stream Segregation 3 4.

Brodbeck C, Jiao A, Hong LE, Simon JZ. 2020b. Neural speech restoration at the cocktail party: Auditory cortex recovers masked speech of both attended and ignored speakers. PLOS Biol. 18:e3000883.

Brodbeck C, Presacco A, Simon JZ. 2018. Neural source dynamics of brain responses to continuous stimuli: Speech processing from acoustics to comprehension. Neuroimage. 172:162–174.

Broderick MP, Anderson AJ, Di Liberto GM, Crosse MJ, Lalor EC. 2018. Electrophysiological Correlates of Semantic Dissimilarity Reflect the Comprehension of Natural, Narrative Speech. Curr Biol. 28:803-809.e3.

Broderick MP, Anderson AJ, Lalor EC. 2019. Semantic Context Enhances the Early Auditory Encoding of Natural Speech. J Neurosci. 0584–19.

Bronkhorst AW. 2015. The cocktail-party problem revisited: early processing and selection of multi-talker speech. Attention, Perception, Psychophys. 77:1465–1487.

Brouwer S, Van Engen KJ, Calandruccio L, Bradlow AR. 2012. Linguistic contributions to speech-on-speech masking for native and non-native listeners: Language familiarity and semantic content. Cit J Acoust Soc Am. 131:1101.

Brungart DS, Simpson BD, Ericson MA, Scott KR. 2001. Informational and energetic masking effects in the perception of multiple simultaneous talkers. J Acoust Soc Am. 110:2527– 2538.

Bryden MP. 1964. The Manipulation of strategies of report in dichotic listening. Can J Psychol. 18:126–138.

Buiatti M, Peña M, Dehaene-Lambertz G. 2009. Investigating the neural correlates of continuous speech computation with frequency-tagged neuroelectric responses. Neuroimage. 44:509–519.

Calandruccio L, Dhar S, Bradlow AR. 2010. Speech-on-speech masking with variable access to the linguistic content of the masker speech. J Acoust Soc Am. 128:860–869.

Carey D, Mercure E, Pizzioli F, Aydelott J. 2014. Auditory semantic processing in dichotic listening: Effects of competing speech, ear of presentation, and sentential bias on N400s to spoken words in context. Neuropsychologia. 65:102–112.

Carlyon RP. 2004. How the brain separates sounds. Trends Cogn Sci. 8:465–471.

Carlyon RP, Cusack R, Foxton JM, Robertson IH. 2001. Effects of attention and unilateral neglect on auditory stream segregation. J Exp Psychol Hum Percept Perform. 27:115–127.

Cherry CE. 1953. Some experiments on the recognition of speech, with one and two ears. J Acoust Soc Am. 25:975–979.

Conway ARA, Cowan N, Bunting MF. 2001. The cocktail party phenomenon revisited: The importance of working memory capacity. Psychon Bull Rev. 8:331–335.

Cooke M. 2006. A glimpsing model of speech perception in noise. J Acoust Soc Am. 119:1562– 1573.

Cooke M, Garcia Lecumberri ML, Barker J. 2008. The foreign language cocktail party problem: Energetic and informational masking effects in non-native speech perception. J Acoust Soc Am. 123:414–427.

Corbetta M, Patel G, Shulman GL. 2008. The reorienting system of the human brain: from environment to theory of mind. Neuron. 58:306–324.

Cowan N, Elliott EM, Scott Saults J, Morey CC, Mattox S, Hismjatullina A, Conway ARA. 2005. On the capacity of attention: Its estimation and its role in working memory and cognitive aptitudes. Cogn Psychol. 51:42–100.

Dale AM, Liu AK, Fischl BR, Buckner RL, Belliveau JW, Lewine JD, Halgren E. 2000. Dynamic statistical parametric mapping: combining fMRI and MEG for high-resolution imaging of cortical activity. Neuron. 26:55–67.

David S V, Mesgarani N, Shamma SA. 2007. Estimating sparse spectro-temporal receptive fields with natural stimuli. Netw Comput Neural Syst. 18:191–212.

Davis MH, Ford MA, Kherif F, Johnsrude IS. 2011. Does semantic context benefit speech understanding through “top-down” processes? evidence from time-resolved sparse fMRI. J Cogn Neurosci. 23:3914–3932.

Davis MH, Johnsrude IS. 2003. Hierarchical processing in spoken language comprehension. J Neurosci. 23:3423–3431.

Deutsch JA, Deutsch D. 1963. Attention: Some theoretical considerations. Psychol Rev. 70:80– 90.

Ding N, Melloni L, Zhang H, Tian X, Poeppel D. 2016. Cortical tracking of hierarchical linguistic structures in connected speech. Nat Neurosci. 19:158–164.

Ding N, Pan X, Luo C, Su N, Zhang W, Zhang J. 2018. Attention Is Required for Knowledge-Based Sequential Grouping: Insights from the Integration of Syllables into Words. J Neurosci. 38:1178–1188.

Ding N, Simon JZ. 2012. Emergence of neural encoding of auditory objects while listening to competing speakers. Proc Natl Acad Sci. 109:11854–11859.

Ding N, Simon JZ. 2014. Cortical entrainment to continuous speech: functional roles and interpretations. Front Hum Neurosci. 8:311.

Donhauser PW, Baillet S. 2019. Two Distinct Neural Timescales for Predictive Speech Processing. Neuron.

Driver J. 2001. A selective review of selective attention research from the past century. Br J Psychol. 92 Part 1:53–78.

Dronkers NF, Wilkins DP, Van Valin RD, Redfern BB, Jaeger JJ. 2004. Lesion analysis of the brain areas involved in language comprehension. Cognition. 92:145–177.

Drullman R, Bronkhorst AW. 2004. Speech perception and talker segregation: effects of level, pitch, and tactile support with multiple simultaneous talkers. J Acoust Soc Am. 116:3090– 3098.

Dupoux E, Kouider S, Mehler J. 2003. Lexical Access Without Attention? Explorations Using Dichotic Priming. J Exp Psychol Hum Percept Perform. 29:172–184.

Durlach NI, Mason CR, Shinn-Cunningham BG, Arbogast TL, Colburn HS, Kidd G. 2003. Informational masking: Counteracting the effects of stimulus uncertainty by decreasing target-masker similarity. J Acoust Soc Am. 114:368–379.

Edin F, Klingberg T, Johansson P, McNab F, Tegnér J, Compte A. 2009. Mechanism for top-down control of working memory capacity. Proc Natl Acad Sci U S A. 106:6802–6807.

Elliott EM, Briganti AM. 2012. Investigating the role of attentional resources in the irrelevant speech effect. Acta Psychol (Amst). 140:64–74.

Fairnie J, Brian BCJ, Remington A. 2016. Missing a Trick: Auditory Load Modulates Conscious Awareness in Audition. J Exp Psychol Hum Percept Perform. 42:930–938.

Fedorenko E, Blank IA. 2020. Broca’s Area Is Not a Natural Kind. Trends Cogn Sci.

Fedorenko E, Scott TL, Brunner P, Coon WG, Pritchett B, Schalk G, Kanwisher N. 2016. Neural correlate of the construction of sentence meaning. Proc Natl Acad Sci U S A. 113:E6256– E6262.

Fiedler L, Wöstmann M, Herbst SK, Obleser J. 2019. Late cortical tracking of ignored speech facilitates neural selectivity in acoustically challenging conditions. Neuroimage. 186:33–42.

Fischl B, Sereno MI, Tootell RBH, Dale AM. 1999. High-Resolution Intersubject Averaging and a Coordinate System for the Cortical Surface, Hum. Brain Mapping.

Fogerty D, Carter BL, Healy EW. 2018. Glimpsing speech in temporally and spectro-temporally modulated noise. J Acoust Soc Am. 143:3047–3057.

Forster S, Lavie N. 2008. Failures to ignore entirely irrelevant distractors: the role of load. J Exp Psychol Appl. 14:73–83.

Francart T, Van Wieringen A, Wouters J. 2010. Comparison of fluctuating maskers for speech recognition tests. Int J Audiol. 50:2–13.

Freyman RL, Balakrishnan U, Helfer KS. 2001. Spatial release from informational masking in speech recognition. J Acoust Soc Am. 109:2112–2122.

Friederici AD, Kotz SA. 2003. The brain basis of syntactic processes: Functional imaging and lesion studies. In: NeuroImage. Academic Press Inc. p. S8–S17.

Gagné JP, Besser J, Lemke U. 2017. Behavioral assessment of listening effort using a dual-task paradigm: A review. Trends Hear. 21:1–25.

Gallun FJ, Diedesch AC. 2013. Exploring the factors predictive of informational masking in a speech recognition task. In: Proceedings of Meetings on Acoustics. Acoustical Society of AmericaASA. p. 060145–060145.

Gazzaley A, Nobre AC. 2012. Top-down modulation: bridging selective attention and working memory. Trends Cogn Sci. 16:129–135.

Getz H, Ding N, Newport EL, Poeppel D. 2018. Cortical tracking of constituent structure in language acquisition. Cognition. 181:135–140.

Glasser MF, Coalson TS, Robinson EC, Hacker CD, Harwell J, Yacoub E, Ugurbil K, Andersson J, Beckmann CF, Jenkinson M, Smith SM, Van Essen DC. 2016. A multi-modal parcellation of human cerebral cortex. Nature. 536:171–178.

Glucksberg S, Cowen GN. 1970. Memory for nonattended auditory material. Cogn Psychol. 1:149–156.

Gramfort A. 2013. MEG and EEG data analysis with MNE-Python. Front Neurosci. 7.

Gramfort A, Luessi M, Larson E, Engemann DA, Strohmeier D, Brodbeck C, Parkkonen L, Hämäläinen MS. 2014. MNE software for processing MEG and EEG data. Neuroimage. 86:446–460.

Hagoort P. 2003. How the brain solves the binding problem for language: A neurocomputational model of syntactic processing. In: NeuroImage. Academic Press Inc. p. S18–S29.

Hahn B, Robinson BM, Leonard CJ, Luck SJ, Gold JM. 2018. Posterior parietal cortex dysfunction is central to working memory storage and broad cognitive deficits in schizophrenia. J Neurosci. 38:8378–8387.

Hervais-Adelman AG, Carlyon RP, Johnsrude IS, Davis MH. 2012. Brain regions recruited for the effortful comprehension of noise-vocoded words. Lang Cogn Process. 27:1145–1166.

Hickok G, Poeppel D. 2007. The cortical organization of speech processing. Nat Rev Neurosci. 8:393–402.

Hill KT, Miller LM. 2010. Auditory attentional control and selection during cocktail party listening. Cereb Cortex. 20:583–590.

Hohlfeld A, Sommer W. 2005. Semantic processing of unattended meaning is modulated by additional task load: Evidence from electrophysiology. Cogn Brain Res. 24:500–512.

Horton C, D’Zmura M, Srinivasan R. 2013. Suppression of competing speech through entrainment of cortical oscillations. J Neurophysiol. 109:3082–3093.

Humphries C, Binder JR, Medler DA, Liebenthal E. 2006. Syntactic and Semantic Modulation of Neural Activity during Auditory Sentence Comprehension. J Cogn Neurosci. 18:665–679.

Iyer N, Brungart DS, Simpson BD. 2010. Effects of target-masker contextual similarity on the multimasker penalty in a three-talker diotic listening task. J Acoust Soc Am. 128:2998– 3010.

Kahneman D. 1973. Attention and effort. Prentice-Hall.

Kidd G, Mason CR, Richards VM, Gallun FJ, Durlach NI. 2008. Informational Masking. Springer, Boston, MA. p. 143–189.

Kidd G, Mason CR, Swaminathan J, Roverud E, Clayton KK, Best V, Best V. 2016. Determining the energetic and informational components of speech-on-speech masking. J Acoust Soc Am. 140:132.

Kilman L, Zekveld A, Hällgren M, Rönnberg J. 2014. The influence of non-native language proficiency on speech perception performance. Front Psychol. 5:651.

Lachter J, Forster KI, Ruthruff E. 2004. Forty-Five Years After Broadbent (1958): Still No Identification Without Attention [WWW Document].

Lambez B, Agmon G, Har-Shai Yahav P, Rassovsky Y, Zion Golumbic E. 2020. Paying attention to speech: The role of working memory capacity and professional experience. Attention, Perception, Psychophys.

Lavie N, Beck DM, Konstantinou N. 2014. Blinded by the load: attention, awareness and the role of perceptual load. Philos Trans R Soc Lond B Biol Sci. 369:20130205.

Lavie N, Hirst A, de Fockert JW, Viding E. 2004. Load Theory of Selective Attention and Cognitive Control. J Exp Psychol Gen. 133:339–354.

Lewis JL. 1970. Semantic processing of unattended messages using dichotic listening. J Exp Psychol.

Linden DEJ. 2007. The working memory networks of the human brain. Neuroscientist.

Luo C, Ding N. 2020. Delta-band Cortical Tracking of Acoustic and Linguistic Features in Natural Spoken Narratives. bioRxiv. 2020.07.31.231431.

Ma L, Steinberg JL, Hasan KM, Narayana PA, Kramer LA, Moeller FG. 2012. Working memory load modulation of parieto-frontal connections: Evidence from dynamic causal modeling. Hum Brain Mapp. 33:1850–1867.

Makov S, Sharon O, Ding N, Ben-Shachar M, Nir Y, Zion Golumbic EM. 2017. Sleep Disrupts High-Level Speech Parsing Despite Significant Basic Auditory Processing. J Neurosci. 37:7772– 7781.

Makov S, Zion Golumbic E. 2020. Irrelevant Predictions: Distractor Rhythmicity Modulates Neural Encoding in Auditory Cortex. Cereb Cortex. 30:5792–5805.

Marrone N, Mason CR, Kidd G. 2008. Tuning in the spatial dimension: Evidence from a masked speech identification task. J Acoust Soc Am. 124:1146–1158.

Matchin W, Hammerly C, Lau E. 2017. The role of the IFG and pSTS in syntactic prediction: Evidence from a parametric study of hierarchical structure in fMRI. Cortex. 88:106–123.

Mattys SL, Davis MH, Bradlow AR, Scott SK. 2012. Speech recognition in adverse conditions: A review. Lang Cogn Process. 27:953–978.

Matusz PJ, Dikker S, Huth AG, Perrodin C. 2019. Are We Ready for Real-world Neuroscience? J Cogn Neurosci. 31:327–338.

McNab F, Klingberg T. 2008. Prefrontal cortex and basal ganglia control access to working memory. Nat Neurosci. 11:103–107.

Mesgarani N, Chang EF. 2012. Selective cortical representation of attended speaker in multi-talker speech perception. Nature. 485:233–236.

Meyer L, Cunitz K, Obleser J, Friederici AD. 2014. Sentence processing and verbal working memory in a white-matter-disconnection patient. Neuropsychologia. 61:190–196.

Meyer L, Grigutsch M, Schmuck N, Gaston P, Friederici AD. 2015. Frontal-posterior theta oscillations reflect memory retrieval during sentence comprehension. Cortex. 71:205–218.

Miller KJ, Sorensen LB, Ojemann JG, Den Nijs M. 2009. Power-law scaling in the brain surface electric potential. PLoS Comput Biol. 5:1000609.

Moray N. 1959. Attention in Dichotic Listening: Affective Cues and the Influence of Instructions. Q J Exp Psychol. 11:56–60.

Morgan EU, van der Meer A, Vulchanova M, Blasi DE, Baggio G. 2020. Meaning before grammar: A review of ERP experiments on the neurodevelopmental origins of semantic processing. Psychon Bull Rev.

Müller JA, Wendt D, Kollmeier B, Debener S, Brand T. 2019. Effect of Speech Rate on Neural Tracking of Speech. Front Psychol. 10:449.

Murphy S, Spence C, Dalton P. 2017. Auditory perceptual load: A review. Hear Res. 352:40–48.

Naveh-Benjamin M, Kilb A, Maddox GB, Thomas J, Fine HC, Chen T, Cowan N. 2014. Older adults do not notice their names: A new twist to a classic attention task. J Exp Psychol Learn Mem Cogn. 40:1540–1550.

Neely C, LeCompte D. 1999. The importance of semantic similarity to the irrelevant speech effect. Mem Cogn. 27:37–44.

Nelson MJ, El Karoui I, Giber K, Yang X, Cohen L, Koopman H, Cash SS, Naccache L, Hale JT, Pallier C, Dehaene S. 2017. Neurophysiological dynamics of phrase-structure building during sentence processing. Proc Natl Acad Sci U S A. 114:E3669–E3678.

Nourski K V, Reale RA, Oya H, Kawasaki H, Kovach CK, Chen H, Howard MA, Brugge JF. 2009. Temporal Envelope of Time-Compressed Speech Represented in the Human Auditory Cortex. J Neurosci. 29:15564–15574.

Nozaradan S, Schönwiesner M, Keller PE, Lenc T, Lehmann A. 2018. Neural bases of rhythmic entrainment in humans: critical transformation between cortical and lower-level representations of auditory rhythm. Eur J Neurosci. 47:321–332.

O’Sullivan JA, Power AJ, Mesgarani N, Rajaram S, Foxe JJ, Shinn-Cunningham BG, Slaney M, Shamma SA, Lalor EC. 2015. Attentional Selection in a Cocktail Party Environment Can Be Decoded from Single-Trial EEG. Cereb Cortex. 25:1697–1706.

Obleser J, Kotz SA. 2010. Expectancy constraints in degraded speech modulate the language comprehension network. Cereb Cortex. 20:633–640.

Olguin A, Bekinschtein TA, Bozic M. 2018. Neural encoding of attended continuous speech under different types of interference. J Cogn Neurosci. 30:1606–1619.

Østby Y, Tamnes CK, Fjell AM, Walhovd KB. 2011. Morphometry and connectivity of the fronto-parietal verbal working memory network in development. Neuropsychologia. 49:3854– 3862.

Oswald CJP, Tremblay S, Jones DM. 2000. Disruption of comprehension by the meaning of irrelevant sound. Memory. 8:345–350.

Parmentier FBR. 2008. Towards a cognitive model of distraction by auditory novelty: The role of involuntary attention capture and semantic processing. Cognition. 109:345–362.

Parmentier FBR, Pacheco-Unguetti AP, Valero S. 2018. Food words distract the hungry: Evidence of involuntary semantic processing of task-irrelevant but biologically-relevant unexpected auditory words. PLoS One. 13:e0190644.

Peelle JE. 2018. Listening effort: How the cognitive consequences of acoustic challenge are reflected in brain and behavior. Ear Hear. 39:204–214.

Postle BR, Ferrarelli F, Hamidi M, Feredoes E, Massimini M, Peterson M, Alexander A, Tononi G. 2006. Repetitive transcranial magnetic stimulation dissociates working memory manipulation from retention functions in the prefrontal, but not posterior parietal, cortex. J Cogn Neurosci. 18:1712–1722.

Pritchard WS. 1992. The brain in fractal time: 1/f-like power spectrum scaling of the human electroencephalogram. Int J Neurosci. 66:119–129.

Pulvermüller F, Shtyrov Y, Hasting AS, Carlyon RP. 2008. Syntax as a reflex: Neurophysiological evidence for early automaticity of grammatical processing. Brain Lang. 104:244–253.

Puvvada KC, Simon JZ. 2017. Cortical Representations of Speech in a Multitalker Auditory Scene. J Neurosci. 37:9189–9196.

Pylkkänen L. 2019. The neural basis of combinatory syntax and semantics. Science (80-).

Rämä P, Relander-Syrjänen K, Carlson S, Salonen O, Kujala T. 2012. Attention and semantic processing during speech: An fMRI study.

Rhebergen KS, Versfeld NJ, Dreschler WA. 2005. Release from informational masking by time reversal of native and non-native interfering speech. J Acoust Soc Am. 118:1274–1277.

Rimmele JM, Zion Golumbic EM, Schröger E, Poeppel D. 2015. The effects of selective attention and speech acoustics on neural speech-tracking in a multi-talker scene. Cortex. 68.

Risko EF, Richardson DC, Kingstone A. 2016. Breaking the Fourth Wall of Cognitive Science. Curr Dir Psychol Sci. 25:70–74.

Rivenez M, Darwin CJ, Guillaume A. 2006. Processing unattended speech. J Acoust Soc Am. 119:4027–4040.

Rivenez M, Guillaume A, Bourgeon L, Darwin CJ. 2008. Effect of voice characteristics on the attended and unattended processing of two concurrent messages. Eur J Cogn Psychol. 20:967–993.

Röer JP, Körner U, Buchner A, Bell R. 2017a. Semantic priming by irrelevant speech. Psychon Bull Rev. 24:1205–1210.

Röer JP, Körner U, Buchner A, Bell R. 2017b. Attentional capture by taboo words: A functional view of auditory distraction. Emotion. 17:740–750.

Rosen S, Souza P, Ekelund C, Majeed AA. 2013. Listening to speech in a background of other talkers: Effects of talker number and noise vocoding. J Acoust Soc Am. 133:2431–2443.

Rottschy C, Langner R, Dogan I, Reetz K, Laird AR, Schulz JB, Fox PT, Eickhoff SB. 2012. Modelling neural correlates of working memory: A coordinate-based meta-analysis. Neuroimage. 60:830–846.

Schell M, Zaccarella E, Friederici AD. 2017. Differential cortical contribution of syntax and semantics: An fMRI study on two-word phrasal processing. Cortex. 96:105–120.

Schepman A, Rodway P, Pritchard H. 2016. Right-lateralized unconscious, but not conscious, processing of affective environmental sounds. Laterality. 21:606–632.

Shavit-Cohen K, Zion Golumbic E. 2019. The Dynamics of Attention Shifts Among Concurrent Speech in a Naturalistic Multi-speaker Virtual Environment. Front Hum Neurosci. 13:386.

Shinn-Cunningham BG. 2008. Object-based auditory and visual attention. Trends Cogn Sci. 12:182–186.

Smirnov D, Glerean E, Lahnakoski JM, Salmi J, Jääskeläinen IP, Sams M, Nummenmaa L. 2014. Fronto-parietal network supports context-dependent speech comprehension. Neuropsychologia. 63:293–303.

Surprenant AM, Neath I, LeCompte DC. 1999. Irrelevant Speech, Phonological Similarity, and Presentation Modality. Memory. 7:405–420.

Swaminathan J, Mason CR, Streeter TM, Best V, Kidd G, Patel AD. 2015. Musical training, individual differences and the cocktail party problem. Sci Rep. 5:1–11.

Teoh ES, Lalor EC. 2019. EEG decoding of the target speaker in a cocktail party scenario: Considerations regarding dynamic switching of talker location. J Neural Eng. 16.

Todd JJ, Marois R. 2004. Capacity limit of visual short-term memory in human posterior parietal cortex. Nature. 428:751–754.

Treisman AM. 1960. Contextual cues in selective listening. Q J Exp Psychol. 12:242–248.

Tun PA, O’Kane G, Wingfield A. 2002. Distraction by competing speech in young and older adult listeners. Psychol Aging. 17:453–467.

Vachon F, Marsh JE, Labonté K. 2019. The Automaticity of Semantic Processing Revisited: Auditory Distraction by a Categorical Deviation. J Exp Psychol Gen. 149.

Vandenberghe R, Nobre AC, Price CJ. 2002. The Response of Left Temporal Cortex to Sentences. J Cogn Neurosci. 14:550–560.

Vandierendonck A. 2014. Symbiosis of executive and selective attention in working memory. Front Hum Neurosci. 8:588.

Vestergaard MD, Fyson NRC, Patterson RD. 2011. The mutual roles of temporal glimpsing and vocal characteristics in cocktail-party listening. J Acoust Soc Am. 130:429–439.

Wild CJ, Yusuf a., Wilson DE, Peelle JE, Davis MH, Johnsrude IS. 2012. Effortful Listening: The Processing of Degraded Speech Depends Critically on Attention. J Neurosci. 32:14010– 14021.

Wood N, Cowan N. 1995. The cocktail party phenomenon revisited: How frequent are attention shifts to one’s name in an irrelevant auditory channel? J Exp Psychol Learn Mem Cogn. 21:255–260.

Yates AJ. 1965. Delayed Auditory Feedback and Shadowing. Q J Exp Psychol. 17:125–131.

Yue Q, Martin RC, Cris Hamilton A, Rose NS. 2019. Non-perceptual Regions in the Left Inferior Parietal Lobe Support Phonological Short-term Memory: Evidence for a Buffer Account? Cereb Cortex. 29:1398–1413.

Zion Golumbic EM, Cogan GB, Schroeder CE, Poeppel D. 2013. Visual input enhances selective speech envelope tracking in auditory cortex at a “Cocktail Party.” J Neurosci. 33.

Zion Golumbic EM, Ding N, Bickel S, Lakatos P, Schevon CA, McKhann G, Goodman RR, Emerson R, Mehta AD, Simon JZ, Poeppel D, Schroeder CE. 2013. Mechanisms Underlying Selective Neuronal Tracking of Attended Speech at a Cocktail Party. Neuron. 77:980–991.

